# Optical Genome and Epigenome Mapping of Clear Cell Renal Cell Carcinoma

**DOI:** 10.1101/2022.10.11.511152

**Authors:** Sapir Margalit, Zuzana Tulpová, Yael Michaeli, Tahir Detinis Zur, Jasline Deek, Sivan Louzoun-Zada, Gil Nifker, Assaf Grunwald, Yuval Scher, Leonie Schütz, Elmar Weinhold, Yehudit Gnatek, Dorit Omer, Benjamin Dekel, Eitan Friedman, Yuval Ebenstein

## Abstract

Cancer cells display complex genomic aberrations that include large-scale genetic rearrangements and epigenetic modulation that are not easily characterized by short-read sequencing. We present a method for simultaneous profiling of long-range genetic/epigenetic changes in matched cancer samples. Clear cell renal cell carcinoma (ccRCC) is the most common subtype of kidney cancer. Most ccRCC cases demonstrate somatic genomic alterations involving the short arm of chromosome 3 (3p), most often targeting the von Hippel–Lindau (*VHL*) gene. Aiming to identify somatic alterations that characterize early stage ccRCC, we performed comprehensive genetic, cytogenetic and epigenetic analyses comparing ccRCC tumor to adjacent non-tumorous tissue. Optical genome mapping identified genomic aberrations such as structural and copy number variations, complementing exome-sequencing results. Single-molecule methylome and hydroxymethylome mapping revealed multiple differential regions, some of them known to be associated with ccRCC pathogenesis. Among them, metabolic pathways were significantly enriched. Moreover, significant global epigenetic differences were detected between the tumor and the adjacent non-tumorous tissue, and a correlation between epigenetic signals and gene expression was found. This is the first reported comparison of a human tumor and a matched tissue by optical genome/epigenome mapping, revealing well-established and novel somatic aberrations.

## INTRODUCTION

Clear cell renal cell carcinoma (ccRCC) is the most common type of renal carcinoma, and its incidence has been increasing in recent years. Over 90% of ccRCC cases demonstrate distinctive changes to the short arm of chromosome 3 (3p), from translocations and deletions to the loss of the entire chromosomal arm. The von Hippel–Lindau (*VHL*) gene, located on this arm, is mutated in 30–56% of sporadic clear cell carcinomas, and is silenced by promoter hypermethylation in up to 19% of cases (1). In these cases, the inactivation of *VHL* has been identified as the earliest event driving the disease. *VHL* loss in ccRCC affects multiple cellular processes as angiogenesis, cell cycle, cell growth, and metabolism (1, 2). However, biallelic *VHL* inactivation alone is not sufficient to induce ccRCC (3, 4). Pathogenic sequence variants affecting other 3p-residing tumor suppressor genes are also frequently observed in ccRCC. Genes such as *PBRM1, SETD2*, and *BAP1*, encode for chromatin and histone modifiers and are often mutated in ccRCC, suggesting a possible role of epigenetic dysregulation in ccRCC tumorigenesis (5). Additionally, DNA copy number variations affecting other chromosomes (e.g., DNA gain and loss in the long arms of chromosomes 5 (5q) and 14 (14q), respectively), are very common in ccRCC (6).

### Epigenetics in ccRCC

One of the fundamental epigenetic mechanisms directly affecting gene expression is DNA methylation of cytosine (5mC) in the dinucleotide sequence CpG. Hypermethylation at promoter and enhancer regions, as well as intragenic hypermethylation, is common in ccRCC, and often results in silencing or inactivation of tumor suppressor genes (7, 8). Moreover, aberrant hypermethylation has been reported to correlate with stage, grade and aggressiveness of RCC, with enhancer hypermethylation being particularly predictive of adverse prognosis (7–9). These and other studies prompted discovery and application of specific prognostic methylation markers in ccRCC (10, 11). In addition to cytosine methylation, 5-hydroxymethylcytosine (5hmC), the oxidation product of 5mC, has gained attention as a modifier of gene regulation, development, and disease. Some suggested mechanisms for the regulatory action of 5hmC include binding to transcription factors, altering chromatin structure through association with histone modifications, modulating alternative splicing *via* binding to related proteins, and involvement in miRNA pathways (12). 5hmC is globally reduced in multiple human cancers (13, 14), including ccRCC (15), and lower 5hmC levels in ccRCC are reportedly associated with poorer prognosis (15, 16). 5hmC data has only recently become available due to the fact that popular methods, such as bisulfite sequencing or methylation arrays, do not distinguish between DNA methylation and hydroxymethylation, and report on their cumulative presence. In order to differentiate these two marks, Tet-Assisted Bisulfite Sequencing (TAB-seq) (17), Oxidative bisulfite sequencing (OxBS-seq) (18), or specific enzymatic labeling such as presented here (14, 19–22) have to be employed.

### Optical genome mapping – the full picture

Optical genome mapping (OGM) in nanochannels is a high-throughput, single-molecule technique that captures ultra-long genomic fragments and may uncover genomic information that is mostly inaccessible by sequencing (23). The generated ultra-long reads make it an appealing method for detecting and validating large-scale genomic rearrangements, such as structural and copy number variations (SVs and CNVs), and repetitive sequence motifs (23, 24). It is considered a superior alternative to cytogenetic analysis and a complementing method to next generation sequencing (NGS). Additionally, unlike NGS methods, where ensemble averages mask cellular variability, OGM provides information at the single cell level, as each mapped DNA molecule originates from a different cell, allowing a high-throughput characterization of cellular heterogeneity (23, 25). Using fluorescence microscopy and designated chemistries, OGM can provide multilayered information from individual DNA molecules. Fluorescent labeling of different genomic features with different colors allows studying multiple epigenetic marks on the single-molecule level, creating a hybrid genetic/epigenetic map for every DNA molecule (19, 23, 24, 26).

Here, we utilize a novel approach to complement single-base resolution exome sequencing with single-molecule optical genome/epigenome mapping. We comprehensively analyze somatic alterations in early stage ccRCC as a demonstration of the ability to apply genome/epigenome mapping to comparative genomics and epigenomics studies.

## MATERIAL AND METHODS

### Patient clinically relevant information

Tumor and normal adjacent tissue were obtained in the course of partial nephrectomy performed in a 66-year-old female. Tumor was diagnosed histologically as clear cell renal cell carcinoma (ccRCC) at pT1a stage (<4 cm in the greatest dimension) with cystic degenerative changes (“early stage ccRCC”). Tumor and normal adjacent tissue were also obtained in the course of radical nephrectomy performed in an 82-year-old male. Tumor was diagnosed histologically as ccRCC with morphological features of eosinophilic variant at pT3a stage (“advanced stage ccRCC”). Tissues were stored from the time of surgery to analysis at −80°C.

Samples collection and handling were approved by institutional review boards in accordance with the declaration of Helsinki.

### Extraction of high molecular weight DNA

Ultra-high molecular weight (UHMW) DNA or high molecular weight (HMW) DNA were extracted using *SP Tissue and Tumor DNA Isolation kit* or *Animal Tissue Isolation kit* according to *Bionano Prep Animal Tissue DNA Isolation Soft Tissue/ Fibrous Tissue Protocol* for normal tissue/ tumor (Bionano Genomics), respectively, according to the manufacturer’s protocol.

### Whole-exome sequencing and analysis

Exome sequencing was provided as a service (CD Genomics). 500 ng DNA were used for library construction. Sequencing libraries were generated using Agilent SureSelect Human All Exon kit (Agilent Technologies) following the manufacturer’s recommendations and index codes were added to attribute sequences to each sample (experimental details provided in the Supplementary Data).

Raw sequencing reads were filtered by Trim Galore software (v0.6.7; 10.5281/zenodo.5127898) to remove reads containing adapters or reads of low quality, so that downstream analyses are based on clean reads. Mapping of paired-end clean reads to the human reference genome (hg38) was performed with Bowtie2 software (v2.2.5; (27)). Following alignment, a pipeline by Genome Analysis Toolkit (GATK, v4.2.2.0; (28)) was followed, including using Samtools (v0.1.19; (29)) for sorting and Picard (https://github.com/broadinstitute/picard/) for marking duplicated reads. Base Quality Recalibration process was applied using standard hg38 reference variants. For variant calling, the alignment files of both samples were firstly merged for efficient simultaneous variant calling. Then, small variants, including single nucleotide polymorphisms (SNPs) and insertions/deletions (InDels) located in exon regions, were called by GATK standard genotype pipeline. The called variants of the different samples were then separated. SnpSift program (v4.3t; (30)) was used to add NCBI dbSNP information (v146; (31)), and SnpEff program (v4.3t; (30)) was used to annotate the variants and determine the effect of each variant. Full pipeline and parameters used can be found in the Supplementary Data.

### DNA barcoding and staining for optical genome mapping

The tumor and adjacent tissue samples were labeled by Direct Label and Stain (DLS) chemistry (DLE-1 enzyme, CTTAAG motive; Bionano Genomics), creating a genetic barcode. Single color labeling was created according to a protocol by Bionano Genomics (https://bionanogenomics.com/support-page/dna-labeling-kit-dls/).

### Dual color labeling for optical epigenome mapping

The early stage ccRCC tumor and adjacent tissue were subjected to two types of epigenetic labeling procedures, to generate a comprehensive optical epigenome map. In order to distinguish the epigenetic marks from the green fluorescent DLE-1 marks, we used the red fluorophore ATTO643 (ATTO-Tech) which was found to perform well under our experimental conditions. Synthetic protocols for the ATTO643 labeling reagents prepared for this study are presented in the Supplementary Data.

#### a. Labeling reduced representation of unmodified cytosines in CpG context (umCpG_rr_)

To create the genetic barcode, 1 μg of U/HMW DNA was mixed with 5X DLE-buffer (to a final concentration of 1X), 2 μL of 20X DL-Green and 2 μL of DLE-1 enzyme (Bionano Genomics) in a total reaction volume of 30 μL for 4 hours at 37°C, immediately followed by heat inactivation at 80°C for 20 minutes. Heat inactivation at these conditions degrades over 97% of the DL-Green cofactor, therefore preventing it from being incorporated by M.TaqI in the following reaction, and making the two reactions orthogonal. Then, unmodified cytosines in the recognition sequence TCGA were fluorescently labeled to perform reduced representation optical methylation mapping (ROM) (24, 32). Two 500 ng reaction tubes of DLE1-labeled DNA were each mixed with 4 μL of 10X CutSmart buffer (New England Biolabs), 60 μM of lab-made synthetic AdoYnATTO643 (see synthesis in the Supplementary Data), 1 μL of M.TaqI (10 units/μL; New England Biolabs) and ultrapure water in a total volume of 40 μL, and incubated for 5 hours at 65°C. Then, 5 μL of Puregene Proteinase K (Qiagen) were added and the reaction tube was incubated for additional 2 hours at 45°C. After the Proteinase K treatment, the two 500 ng reaction tubes were merged and drop-dialyzed as one against 20 mL of 1X TE buffer (pH 8) with 0.1 μm dialysis membrane for a total of 2 hours. Finally, 300 ng recovered dual-color DNA were stained to visualize DNA backbone by mixing it with 15 μL of 4X Flow Buffer (Bionano Genomics), 6 μL of 1M DTT (Bionano Genomics), 3 μL of 0.5M Tris (pH 8), 3 μL of 0.5M NaCl, 4.8 μL of DNA Stain (Bionano Genomics) and ultrapure water to a total volume of 60 μL, and incubated overnight at 4°C. The orthogonality of the two consecutive reactions was confirmed by no observed increase in false DLE-1 labels.

#### b. Labeling 5hmC sites

To create the genetic barcode, 580-750 ng of U/HMW DNA in two reaction tubes were each mixed with 5X DLE-buffer (to a final concentration of 1X), 1.5 μL of 20X DL-Green and 1.5 μL of DLE-1 enzyme (Bionano Genomics) in a total reaction volume of 30-35 μL. The reaction was incubated for 4 hours at 37°C. Then, 5hmC sites were labeled by the enzyme β-glucosyltransferase from T4 phage (T4-BGT) (19). Magnesium chloride was added to 30 μL of DLE-labeled DNA to a final concentration of 9 mM. Then, the DNA was added to 4.5 μL of 10X NEBuffer 4 (New England Biolabs), uridine diphosphate-6-azideglucose (UDP-6-N3-Glu; lab-made, (21)) in a final concentration of 50 μM, 30 units of T4 β-glucosyltransferase (New England Biolabs) and ultra-pure water in a final volume of 45 μL. The reaction mixture was incubated overnight at 37°C. The following day, dibenzocyclooctyl (DBCO)-ATTO643 (see synthesis in the Supplementary Data) was added to a final concentration of 150 μM and the reaction was incubated again at 37°C overnight. The next day, the reaction tubes were added 5 μL of PureGene Proteinase K (Qiagen) and incubated for additional 30 minutes at 50°C. After the Proteinase K treatment, the two identical reaction tubes were merged and drop-dialyzed as one against 20 mL of 1X TE buffer (pH 8) with 0.1 μm dialysis membrane for a total of 2-2.5 hours. Finally, 300 ng recovered dual-color DNA was stained to visualize DNA backbone, by mixing it with 4X Flow Buffer (Bionano Genomics) to a final concentration of 1X, 1M DTT (Bionano Genomics) to a final concentration of 0.1 M, Tris (pH 8) to a concentration of 25 mM, NaCl, to a concentration of 25 mM, EDTA to a final concentration of 0.008-0.01 M, DNA Stain (Bionano Genomics) to a final vol/vol ratio of 8%, and ultrapure water. The reaction mixture was shaken horizontally on a HulaMixer for an hour and then incubated overnight at 4°C.

### Optical mapping

Single molecule data of all samples were generated on a Saphyr instrument (Bionano Genomics) with Saphyr chips (G1.2).

### Structural variant and copy number variation calling

*De novo* assemblies of single-color data were generated by Bionano Access (v1.6.1) with Bionano Solve (v3.6.1). The parameters set were ‘haplotype with extend and split’ and ‘cut CMPR’. The *in silico* digested human genome GRCh38.p13 (*hg38_DLE1_0kb_0labels.cmap*) was used as the reference.

Variants were called using Variant Annotation Pipeline (VAP), performed both as single sample analysis and as dual samples analysis (tumor vs. matched normal tissue) in Bionano Access combined with Bionano Tools (v1.6.1) and Bionano Solve (v3.6.1), with default filters. Only SVs that are not present in the Bionano controls dataset were considered. Copy Number Variation (CNV) analysis was performed using the same tools. CNV allows the detection of large, unbalanced aberrations based on normalized molecule coverage and was performed with default parameters as a part of the *de novo* assembly.

### Optical epigenome mapping analysis

Optical mapping data for each sample were merged to a single dataset using Bionano Access (v1.6.1) and Bionano Solve (v3.6.1). Genetic and epigenetic channels were swapped in these files with Bionano Solve before the alignment of the molecules files to the reference, as instructed by the company. Molecules spanning over 150 kbp were then aligned to the *in silico* human genome reference GRCh38.p13, based on DLE-1 recognition sites (*hg38_DLE1_0kb_0labels.cmap*), with default parameters matching the following combination of arguments: haplotype, human, DLE-1, Saphyr. Only molecules with an alignment confidence equal to or higher than 15 (P <= 10^-15^) that at least 60% of their length was aligned to the reference were used for downstream analysis. Alignment outputs were converted to global epigenetic profiles (bedgraph files) and to single-molecule level epigenetic maps according to the pipeline described by Gabrieli et al. and Sharim et al. (19, 24). For more information, see ebensteinLab/Irys-data-analysis on Github. Only regions covered by at least 20 molecules were considered. The average epigenetic score in each genomic position was calculated as the number of detected epigenetic labels in the position divided by the total number of molecules covering the position. The number of epigenetic labels per 100 kbp in an experiment was calculated as the total number of labels in mapped and filtered reads divided by the total corrected length of the mapped and filtered reads. hg38 genome coverage was calculated for a genome size of 3.1 Gbp. Positions of sequence motifs in the reference were obtained using the R package *BSgenome* (https://bioconductor.org/packages/release/bioc/html/BSgenome.html).

### Definition of annotated genomic regions

Gene bodies were defined as spanning from the transcription start site (TSS) to the transcription end site (TES) annotated by GENCODE v34 (33). Promoters were defined as ranging from 1000 bp upstream to 500 bp downstream from the GENCODE TSS. General predicted enhancers were mapped to gene targets by JEME and adapted from Cao et al. (34). Genomic coordinates of enhancers were converted from the human genome build hg19 to hg38 using UCSC liftOver (35). Enhancers overlapping ambiguous genomic regions (36) were discarded, as well as pairs of enhancers and gene targets that are overlapping or in close proximity (up to 5 kbp). ccRCC-related enhancers were adapted from Yao et al. (37) based on differential H3K27ac and H3K4me1 scores not overlapping with promoters in histone chromatin immunoprecipitation sequencing (ChIP-seq) of 10 primary tumor/ normal pairs, 5 patient-matched tumor-derived cell lines, 2 commercially available ccRCC lines (786-O and A-498), and 2 normal kidney cell lines (HK2 and PCS-400). Some of the enhancers were assigned to target genes. We adapted assignments made by correlations between H3K27ac signals and expression of genes within the same topologically associating domain (TAD), and by a capture-C experiment in 786-O cells. Genomic coordinates of enhancers were converted from the human genome build hg19 to hg38 using UCSC liftOver. ccRCC-related ‘super-enhancers’, regions comprising dense clusters of enhancers located near known regulators of cell identity and disease, were also adapted from Yao et al. and converted to hg38 coordinates. Non-overlapping genomic windows of 1 kbp and 50 kbp of hg38 were generated using Bedtools *makewindow* (v2.26.0; (38)).

### Epigenetic ideograms

The weighted mean of epigenetic signals in 50 kbp genomic windows was calculated using Bedops *bedmap* (v2.4.35, (39)). Ideograms displaying the density of epigenetic labels were created with the R package Rideogram (40) with a minor modification: the values were scaled between 1 and the maximal value in the dataset, times 10,000. The darkest color in a pair of whole-genome ideograms was determined according to the highest value in the adjacent and tumor samples.

### Gene expression data

Publicly available RNA-seq data of three tumor-matched pairs of ccRCC (stage 3) patients (PRJNA396588, GEO accessions: pair 1: GSM2723919, GSM2723920; pair 2: GSM2723927, GSM2723928; pair 3:GSM2723929, GSM2723930; (37)) were aligned to the human genome (hg38) using TopHat (v2.1.0, (41)) with default parameters and library-type and fr-firststrand flags, after retrieving the raw files with NCBI SRA toolkit (42). Only uniquely mapped reads were analyzed (minimal mapping quality of 30). Gene counts were obtained using HTSeq (htseq-count, v0.11.3, (43)) against the GENCODE v34 (33) reference gene models. Transcripts per million (TPM) scores were calculated.

### Epigenetic signals along aggregated genes and enhancers

Transcription start and end sites (TSS and TES) of protein-coding genes were defined according to GENCODE annotation (v34, (33)). Protein-coding genes were divided into four groups based on their average normalized TPM score in the RNA-seq of three tumor-matched ccRCC pairs. Unexpressed genes were defined as genes with TPM value <= 0.01 (~3000 genes). The other expression groups are three equal quantiles of the expressed protein-coding genes (~6000 genes per group). Mean epigenetic signals along genes were calculated using DeepTools *computeMatrix* (v3.4.1, (44)) in scale-regions mode, where each gene was scaled to 15 kbp and divided into 300 bp bins.

Mean epigenetic signals along ccRCC-related enhancers were calculated using DeepTools *computeMatrix* (v3.4.1, (44)) in reference-point mode, where the midpoint of the enhancer is the reference point. The midpoint was extended by 9 kbp up and downstream, and the region was divided into 300 bp bins.

Compressed matrix output was summarized by DeepTools *plotProfile*.

### Finding differentially modified regions

The number of epigenetic labels along annotated genomic regions (gene bodies, promoters, general predicted enhancers, ccRCC-related enhancers and super-enhancers; see *Definition of annotated genomic regions*) and 1 kbp genomic windows in individual molecules was counted. Only regions entirely covered by over 20 molecules were regarded. To find differentially modified regions, an independent two-sample t-test was applied to the pool of independent molecules fully covering a region in each condition. Regions with a zero in the t-test’s denominator (when the estimated standard deviations in both samples were zero) where discarded from the analysis.

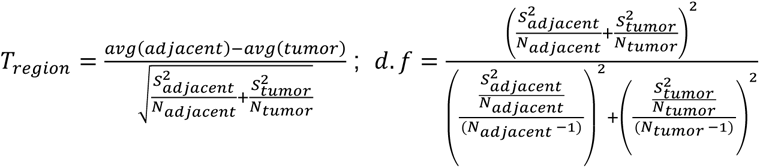

*Equation 1. T-statistics and degrees of freedom (d.f). avg(adjacent*) and *avg(tumor*) are the average number of labels in the region of molecules covering it, in each of the samples. *S_adjacent_ and S_tumor_ are the estimates for the standard deviations in each sample. N_adjacent_ and N_tumor_ are the number of molecules covering the region in each of the samples*.

A p-value was then calculated and a Benjamini-Hochberg false discovery rate (FDR, 1995 (45)) correction was applied. Regions with q<0.1 were considered differentially modified.

### Calculating fold change of epigenetic signals between adjacent and tumor samples

A continuous genome-wide track of fold change (ratio between the epigenetic signal in the adjacent tissue and the tumor) was calculated as follows: bedgraph files containing epigenetic signals in both samples were combined for direct comparison using Bedtools *unionbedg* (v2.26.0, (38)). A pseudo signal of 0.01 was added to each position in either sample to avoid division by 0. Then, the signal in the adjacent tissue was divided by the signal in the tumor to generate the fold change track.

Fold change in discrete 1 kbp genomic windows was calculated as follows: The average number of epigenetic labels in molecules covering the window in each sample was calculated (only windows covered by at least 20 molecules were considered). A pseudo signal of 0.005 was added to each window in either sample to avoid division by 0. Then, the signal in the adjacent tissue was divided by the signal in the tumor. Log2 of this ratio was then calculated.

### Enrichment analysis

The Database for Annotation, Visualization and Integrated Discovery (DAVID; (46, 47)) was used to locate biological terms and pathways that are significantly enriched among unique genes associated with the differentially modified regions. Unique genes associated with all regions covered in the experiment served as background lists. Fisher exact test, corrected by Benjamini-Hochberg false discovery rate (45), was employed to define enriched biological terms and pathways (q<0.05) in ‘functional annotation chart’ analysis.

## RESULTS

We present the first reported comparison of a human tumor and a matched tissue by optical genome/epigenome mapping, revealing disease-relevant and differential SVs, CNVs and epigenetic modifications in these samples. A pathology-classified ccRCC tumor and an adjacent normal kidney tissue (Figure 1.A.), were sequenced for detection of genetic disease signatures (Figure 1.B.). Then, OGM of both samples was performed on the Bionano Genomics Saphyr instrument to provide next-generation cytogenetics (Figure 1.C.). Finally, to obtain optical epigenome mapping, custom labeling chemistries were employed on top of the genetic barcode labels (Figure 1.D.). The dually labeled DNA molecules were then imaged and processed (Figure 1.E.), and the epigenetic signals were inspected on a genome level, locus-specific level and single-molecule level (Figure 1.F.).

**Figure. 1.**
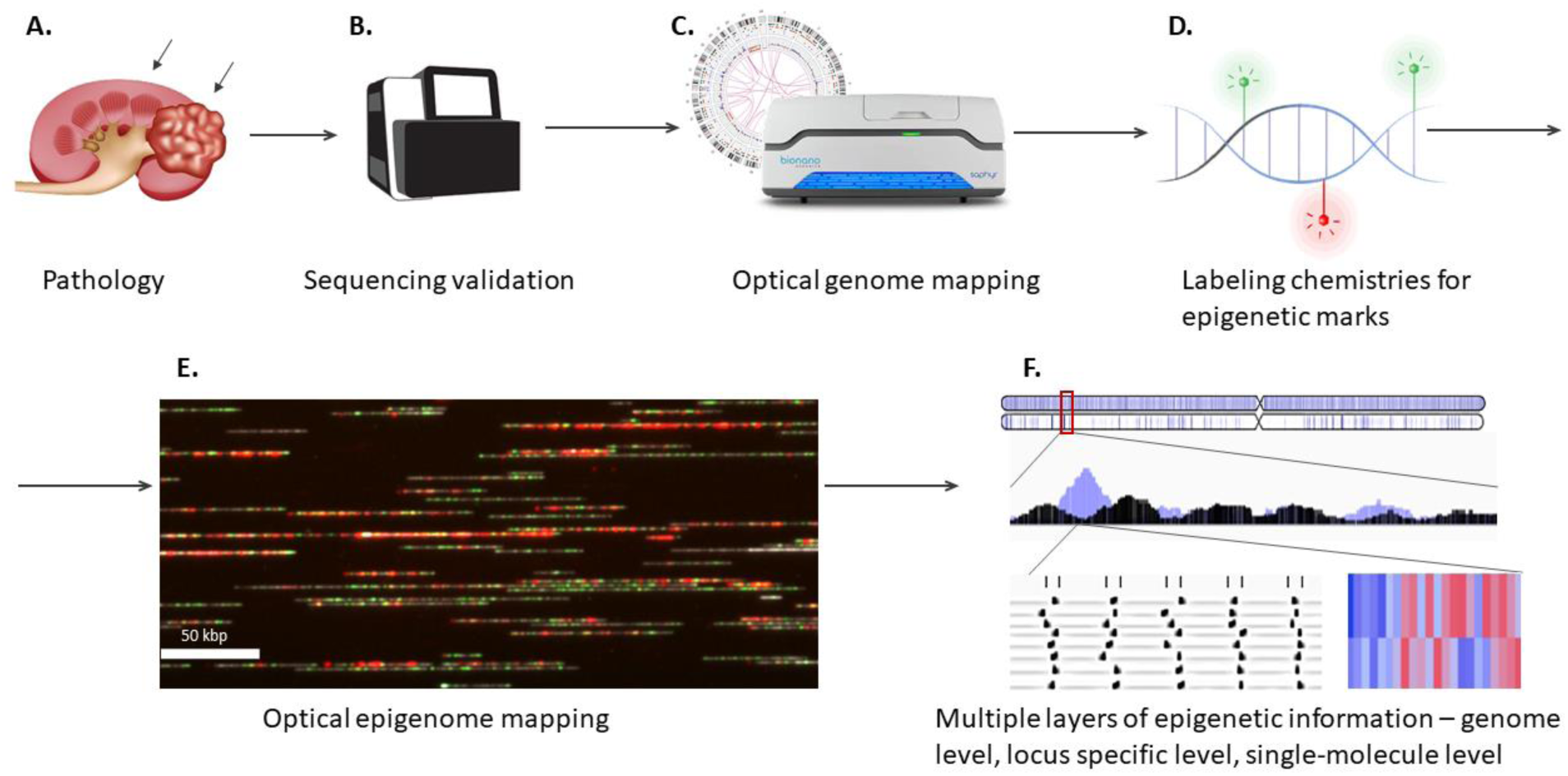
Assay workflow. **A)** A ccRCC tumor and a normal adjacent tissue were pathologically classified and isolated. **B)** Whole-exome of both tissues was sequenced. **C)** Optical genome mapping was applied to detect SVs and CNVs. **D)** Additional labeling schemes for epigenetic features, developed in our lab, were applied to the long DNA molecules from both samples. **E)** Labeled DNA molecules were imaged in three colors (genetic barcode, epigenetic marks, and molecule contour) for optical epigenome mapping. **F)** Multiple levels of epigenetic information were extracted – genome view, chromosome level, locus-specific average, and single-molecule level, in coding and non-coding genomic regions.

### Optical genome mapping detects SVs and CNVs of ccRCC tumor and adjacent tissue

Initial exome-sequencing revealed several genetic aberrations in genes associated with ccRCC (7, 11, 37), including variants with a well-accepted and proven clinical impact in both the early and the advanced stage ccRCC tumors. Noteworthy are variants in *VHL* and *PBRM1* that are known to be highly associated with ccRCC (7, 11, 37). More details about SNPs and InDels discovered by this analysis can be found in the Supplementary Data, in Supplementary Figure S1 and in Supplementary Table S1.

To further investigate the genetic structure of these ccRCC samples, next-generation cytogenetics was employed using OGM. Genetic single-molecule data of early and advanced stage ccRCC tumors and adjacent tissues were generated. An average of 744.3 Gbp (± 52.9 Gbp) of size-filtered (>150 kbp) single-molecule data was generated per sample, with an average molecule N50 of 270.8 kb (± 24.5 kbp). The single-molecule data served to construct an annotated and phased *de novo* assembly for each sample. The average N50 of contigs in all assemblies was 59.1 Mbp (±0.7 Mbp) (full details can be found in Supplementary Table S2).

Bionano Genomics’ Variant Annotation Pipeline (VAP) for normal-tumor pairs revealed 5,666 SVs in the early stage tumor sample and 5,658 SVs in its adjacent normal tissue, 5,842 SVs in the advanced stage tumor, and 5,829 SVs in its adjacent normal tissue. These SVs include insertions, deletions, inversions and duplications, and the distribution between the different types in all samples is similar. The exception is translocation breakpoints, that were detected only in the advanced stage ccRCC tumor. Full details are given in Supplementary Table S3 and Supplementary Figure S3. The detected SVs were compared to Bionano genomics’ healthy controls database to define rare, possibly pathogenic, SVs. Less than 1% of SVs were not found in the healthy database and defined as rare variants. Figure 2.A. shows the rare SVs detected in the four samples in chromosomes 3 and 5. Most (60%-70%) of the rare SVs detected co-localize with a gene or several genes. Among genes overlapping SVs, the advanced stage samples display a dramatic increase in genes overlapping tumor SVs only (63% of genes compared to only 25% in the early stage; Figure 2.B., Supplementary Data and Supplementary Table S4). The *VHL* gene locus did not overlap any rare SV; however, copy number variation (CNV) analysis revealed that one copy of the entire 3p chromosomal arm was lost in the advanced stage tumor (Figure 2.C. and Supplementary Figure S4). In addition to *VHL*, this aneuploidy covers the genes *BAP1, PBRM1* and *SETD2*, that are known to be associated with ccRCC (5, 48). Also deleted is the entire 3p26.3 cytoband, which is known to be associated with deletions causing the 3p-deletion syndrome (Del3p), typically characterized by renal and gastrointestinal abnormalities, in addition to growth retardation and developmental delay (49). Additional aneuploidies, in chromosomes 9, 14 and 5q were detected in this sample (Figure 2.C.), in line with previously reported data for ccRCC. Jonasch and co-workers (48) considered 5q gain to be an alternative ccRCC tumor initiator. Losses of 9p and 14q are considered lethal events followed by metastases, as these chromosomal arms involve genes essential for cell cycle or the metabolism of the *VHL* product, including *CDKN2A* (cyclin-dependent kinase inhibitor 2A) on chromosome 9p (50) or *HIF1A* (hypoxia-inducible factor 1A) on chromosome 14q (5, 48, 51).

**Figure 2.**
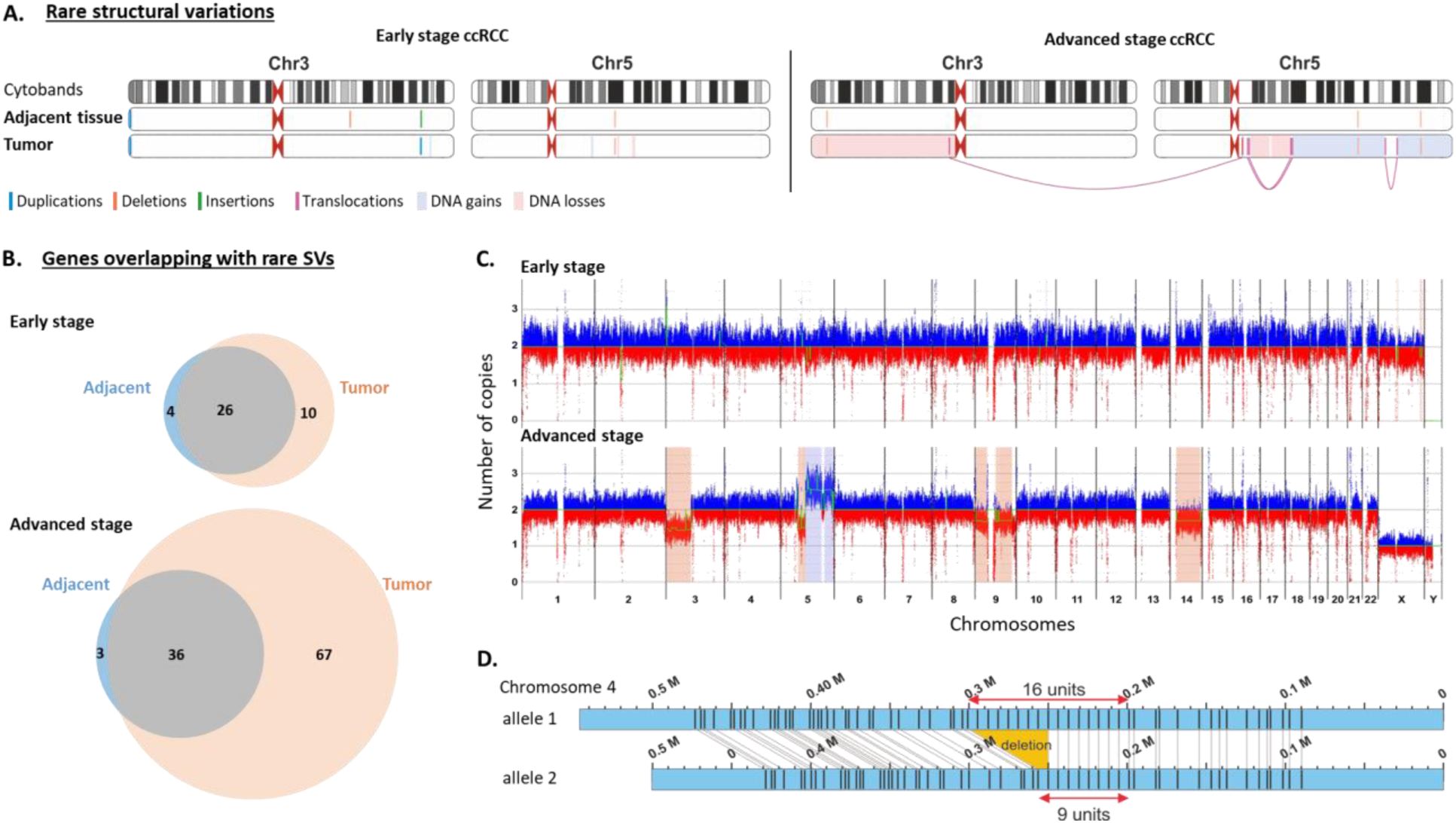
Next-generation cytogenetics with OGM. **A)** Chromosomes 3 and 5 in early and advanced stage ccRCC tumors and normal adjacent tissues. Marked below the cytobands, are rare, possibly pathogenic structural variants. **B)** Venn diagrams depicting genes overlapping with rare structural variations discovered in the early and advanced stage ccRCC tumor and normal adjacent tissues. **C)** Copy number variation profiles of early and advanced stage ccRCC tumors with a baseline set to two copies. DNA gains are colored in blue, DNA losses are colored in red. Aneuploidies are highlighted with colored boxes in the same color scale as DNA gains and losses. **D)** An example of a repeat array with a different number of repeats in two alleles, discovered by OGM, showing a phased chromosomal segment (Conting IDs: 2281, 2282).

Although no such aneuploidies were detected in the early stage samples, several smaller DNA gains and losses were observed, including a DNA gain (segmental duplication) on chromosome 3 spanning the genes *LINC01266* and *CNTN6*. Seven other regions, on chromosomes 3, 5, 10 and X, were found to have DNA gains or losses unique to the tumor sample, ranging between 0.5-6.9 Mb in length (Supplementary Table S5). These CNVs are not known to be associated with ccRCC (48, 52). A CNV on chromosome 10 overlaps the region encoding microRNA-584, which was shown to have significantly lower expression levels in ccRCC tumors, as well as lower cell viability and motility, and was therefore marked as a tumor suppressor microRNA in ccRCC (53). A 1.2 Mb long DNA gain on chromosome 5 with a fractional copy number variation of about 2.4 copies, covers the entire *FOXD1* gene locus. *FOXD1* encodes a forkhead transcription factor belonging to a family of proteins that act as terminal effectors of several key signaling pathways, such as the MAPK pathway. They contribute to the regulation of homeostasis, and their misregulation can induce human genetic diseases including cancer (54). The most extended DNA loss in the early stage samples, on chromosome X, spans about 6.9 Mb in length and contains mainly genes underlying microRNAs or piRNAs.

Several putative events of gene fusion, that can potentially form chimeric genes from the concatenation of independent genes as a byproduct of genomic instability, were detected in both sample pairs (see Supplementary Data). One of them is a fusion of the PCDHA gene cluster, involving 14 genes (*PCDHA1-13, PCDHAC1*), and *AC011346*, that was caused by an intrachromosomal translocation on chromosome 5 (Figure 2.A.).

An important benefit of OGM is the ability to phase structurally-complex regions, such as large repetitive arrays. An example in Figure 2.D. shows allele-specific copy number of a DNA repeat array in chromosome 4.

### Genome-wide and locus-specific epigenetic profiles of unmodified CpGs and 5hmC

Two epigenetic marks were labeled and optically mapped by optical epigenome mapping. 5hmC was directly labeled with a fluorophore according to Gabrieli et al. (19). Unmodified CpG sites within TCGA sequence motifs were labeled according to Sharim et al. (24), to produce a reduced representation map of unmodified CpGs (umCpG_rr_). The obtained map is complementary to a reduced representation map of modified CpG sites. Since most CpG sites in the human genome are methylated (55), labeling the unmodified sites provides increased sensitivity.

For the early stage ccRCC tumor sample, an effective coverage of 73X of the reference genome was collected in 5hmC data and 99X in umCpG_rr_ data. For its normal adjacent tissue, an effective coverage of 48X of the reference genome was collected in 5hmC data and 84X in umCpG_rr_ data. Molecules N50 was ~200 kbp. Full details are provided in Supplementary Table S6.

Figure 3 and Supplementary Figure S5 show a chromosomal distribution and genome-wide global levels of umCpG_rr_ and 5hmC for the early stage ccRCC tumor and the normal adjacent tissue. Most notably, we observed that the 5hmC profile in both tissue samples is sparse, as expected, and a global reduction is observed in the tumor sample compared to the normal adjacent tissue (Figures 3.A. and 3.B. and Supplementary Figure S5.A.), in accordance with global 5hmC reduction known in many types of cancers (15). The umCpG_rr_ signal in the tumor sample is also globally reduced (Figures 3.A. and 3.B. and Supplementary Figure S5.B.), indicating its hypermethylation, in agreement with the literature (8). As expected, the umCpG_rr_ profile is denser than the 5hmC profile in both samples.

**Figure 3.**
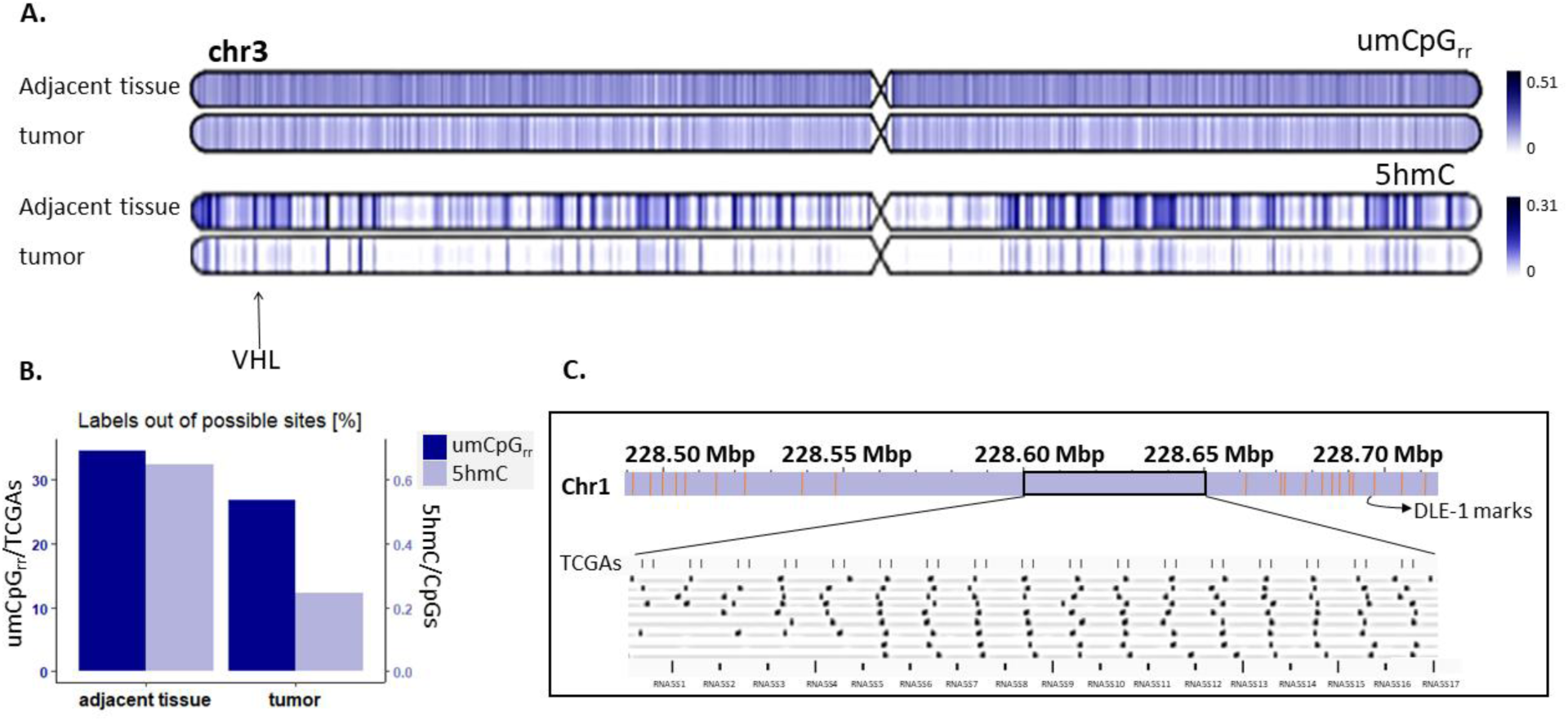
Genome-wide umCpG_rr_ and 5-hmC profiles. **A)** Double Ideograms showing the density of umCpG_rr_ (top) and 5hmC (bottom) along chromosome 3 in the normal adjacent tissue and the early stage ccRCC tumor. **B)** Percentages of detected unmodified CpG sites in TCGA sequence context (umCpG_rr_) out of all appearances of TCGA sites in hg38, and of 5hmC sites detected out of all CpG sites in hg38, in the normal adjacent tissue and the early stage ccRCC tumor. **C)** An example of a repeat array in chromosome 1, not marked by the genetic barcode, but marked with umCpG_rr_ sites. The array corresponds to RNA5S genes.

Epigenetic labeling can also provide complementary genetic information as it addresses regions where the genetic barcode is sparse. This combined approach provides basic genetic information unavailable through other methods, such as repetitive regions. Repetitive elements in DNA are further differentiated by the methylation state of the repeat units, which can affect the function of individual units or even the activity of the entire array. Methylation levels of known repeat arrays were shown to correlate with disease (56–58). An interesting example is a tandem repeat array in chromosome 1 (1.q42), containing a domain of RNA5S genes, encoding 5S rRNA, composed of 17 units according to the human reference, but the actual cluster size is known to vary among individuals (59). Since the repetitive region lacks DLE-1 motifs, the optical genome map cannot provide an accurate copy number of the repeats. Figure 3.C. shows that there are 17 double umCpG_rr_ recognition sites (TCGA) throughout the array in hg38, and that the presence of these sites in the repetitive units allows both counting the copy number and assessing the methylation state of each unit, simultaneously. Supplementary Figure S6 shows fluorescence microscopy images of two representative DNA molecules from the digitized pileup shown in Figure 3.C.

### 5hmC and umCpG_rr_ levels correlate with gene expression

An attractive feature of epigenetic mapping is the ability to relate the epigenetic status of genes to gene expression. To examine this aspect, genes were divided into four groups based on the average TPM value of three ccRCC kidneys in an RNA-seq experiment by Yao et al. ((37); see methods). The mean umCpG_rr_ and 5hmC signals in the adjacent tissue and tumor in each expression group were plotted against the normalized distance from the TSS (Figure 4. A.,B.,D.,E.). In all expression groups, umCpG_rr_ signal around the TSS is lower in the tumor than in the adjacent tissue, and the signal increases with gene expression for both samples (Figure 4.C). In all expression groups, the 5hmC level in gene bodies was lower in the tumor than in the adjacent tissue, and increased with gene expression in both samples (Figure 4.F.). Such correlations with gene expression suggest that epigenetic signals on DNA may serve as a proxy for gene expression, reducing the need to quantify RNA levels. We observed a similar correlation between gene expression and umCpG_rr_ signal in genes for the GM12878 cell line, also supported by whole-genome bisulfite sequencing signal (Supplementary Figure S7). A similar correlation between gene expression and 5hmC signal in genes was previously observed in human peripheral blood cells (19). umCpG_rr_ and 5hmC signals along ccRCC-related enhancers in the early stage ccRCC tumor and its normal adjacent tissue, showing a reduction of both signals in the tumor sample, are presented in Supplementary Figure S8.

**Figure 4.**
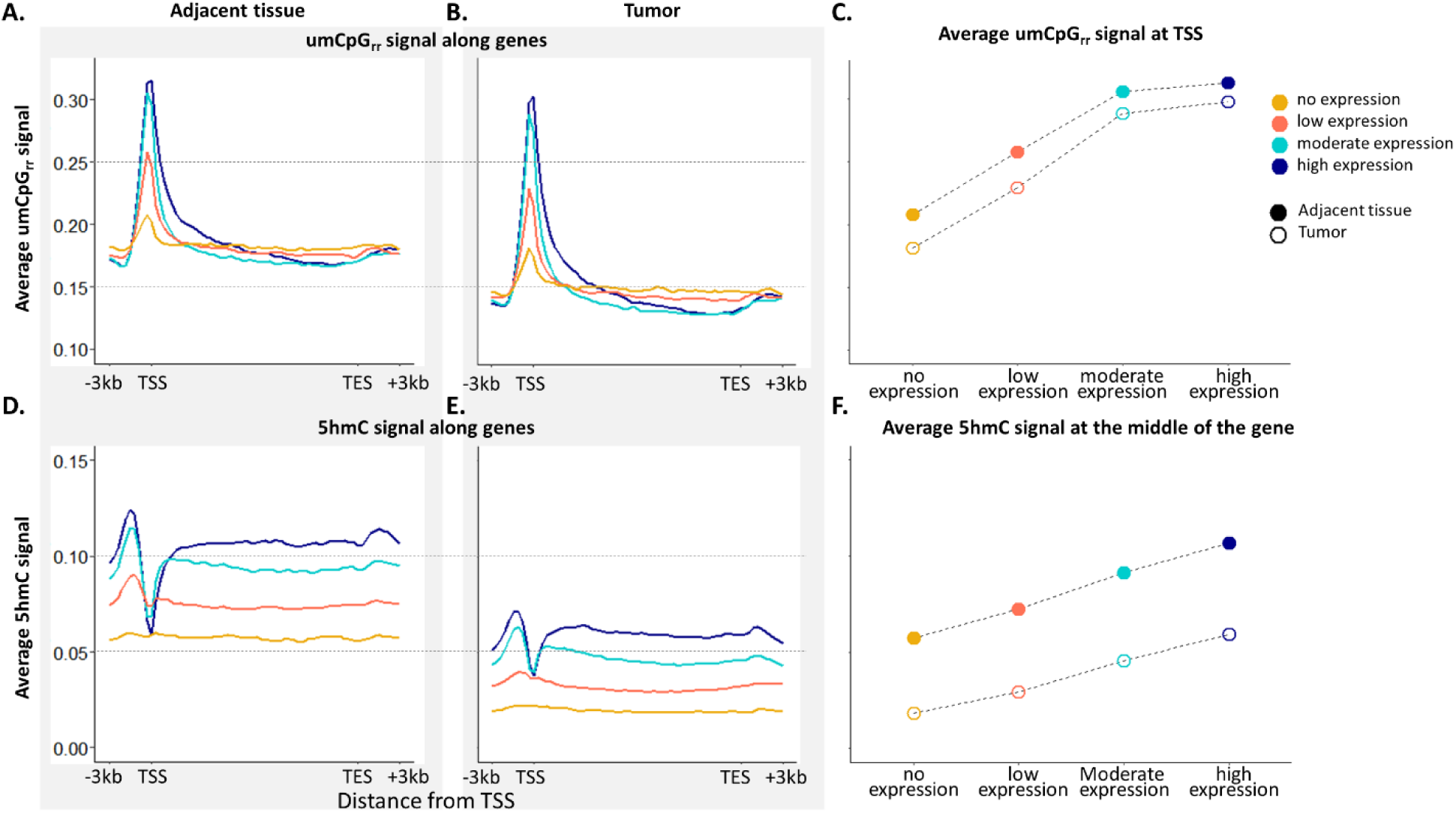
Epigenetic signals along genes as a function of gene expression. **A)** umCpG_rr_-signal in a normal adjacent kidney tissue, four gene groups divided by their expression in adjacent kidney tissues. **B)** umCpG_rr_ signal in a ccRCC tumor, four gene groups divided by their expression in ccRCC tumors. **C)** Average umCpG_rr_ signal at the TSS. **D)**5hmC signal in a normal adjacent kidney tissue, four gene groups divided by their expression in adjacent kidney tissues. **E)**5hmC signal in a ccRCC tumor, four gene groups divided by their expression in ccRCC tumors. **F)** Average 5hmC signal at gene-body midpoints.

### Differentially modified regions

We characterized the differential epigenetic loci in the genomes of the tumor and the adjacent tissue and looked for epigenetically enriched physiological pathways affected by these changes. We applied a differential modification analysis on gene bodies, promoters, general predicted enhancers, ccRCC-related enhancers and ccRCC-related super-enhancers. The genes associated with the differentially modified elements were tested for enrichment of biological terms and pathways.

Several ccRCC-related genes or their associated elements were found to be differentially modified. For example, gene bodies with differential hydroxymethylation include *PRCC*, *BAP1* and *VHL* (displayed in Figure 5.A.). Gene bodies with differential umCpG_rr_ signals include *PBRM1*. General predicted enhancers with differential hydroxymethylation target genes including *PRCC, VEGFA, BAP1* and *VHL* (displayed in Figure 5.A.). General predicted enhancers with differential umCpG_rr_ signals target genes including *VHL* (displayed in Figure 5.A.), *PRCC, BAP1* and *PBRM1*. ccRCC-related enhancers with differential 5hmC and umCpG_rr_ signals target *VEGFA*. Lists of differentially modified elements can be found in Supplementary Tables S7 and S8.

**Figure 5.**
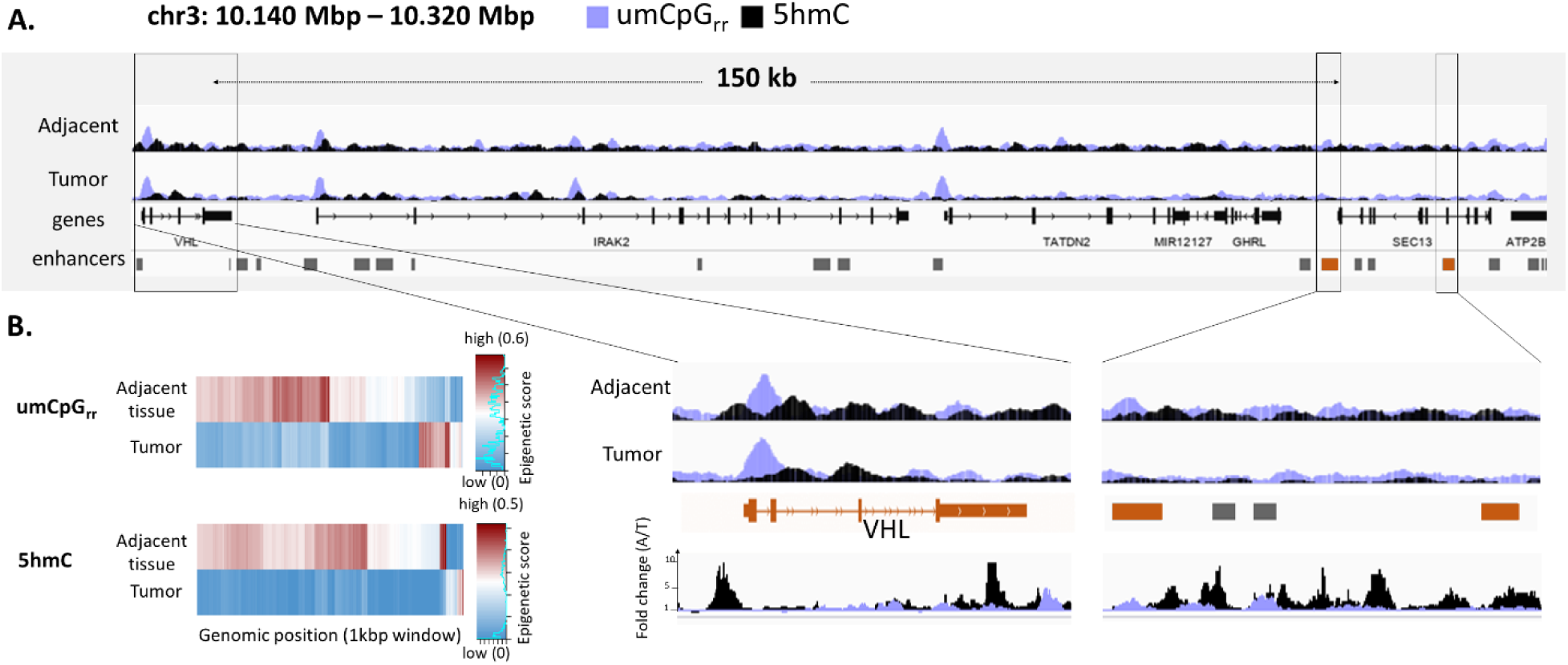
Differentially modified regions. **A)** Integrative Genomics viewer (IGV, (60)) plots of umCpG_rr_ (violet) and 5hmC (black) signals in the adjacent tissue and the ccRCC tumor, along a 180 kbp region in chromosome 3 that covers the *VHL* gene and two general predicted enhancers targeting it. Below is a zoomed-in view of the *VHL* gene and the enhancers. The bottom panel shows plots displaying the fold change of the signals, calculated as the signal in the normal adjacent tissue divided by the signal in the tumor tissue. **B)** Heatmaps of the 250 most differentially umCpG_rr_-modified (top) and 5hmC-modified (bottom) 1 kbp genomic windows between the normal adjacent and the early stage tumor tissue.

Out of 1,114,617 1 kbp genomic windows that contain at least one umCpG_rr_ recognition site and are covered in both samples by at least 20 molecules, 18,489 had differential umCpG_rr_ signal (q<0.1; Supplementary Figure S9.A). These windows overlap 5,190 genes, 87 promoters, 712 general predicted enhancers, 659 ccRCC-related enhancers and 317 ccRCC-related super-enhancers. For 5hmC, 2,744,450 genomic windows containing at least one CpG site were found, but only 537 had significant differential 5hmC signal (q<0.1; Supplementary Figure S9.B). These windows overlap 314 genes, 5 promoters, 22 general predicted enhancers, 31 ccRCC-related enhancers and 19 ccRCC-related super-enhancers. Figure 5.B. presents heatmaps of the 250 most differential 1 kbp genomic windows between the normal adjacent tissue and the early stage tumor.

Alterations in metabolites and dysregulated metabolism are known hallmarks of cancer, and changes in several groups of essential metabolites are manifested in ccRCC. One characteristic of ccRCC cells is that they appear morphologically burdened by lipid and glycogen, suggesting altered fatty acid and glucose metabolism. Additionally, changes in metabolites are associated with tumor progression and metastasis (61). In our case, KEGG pathways enriched among genes that have differential umCpG_rr_ signals are *pentose and glucuronate interconversions* (q=0.0018), *ascorbate and aldarate metabolism* (q=0.071), and *metabolic pathways* (q=0.071). *Metabolic pathways* were also found to be enriched among differentially hydroxymethylated genes (q<10^-10^) and among genes predicted to be targeted by general enhancers with differential hydroxymethylation (q<10^-3^) (Table 1; a full list of enriched biological terms can be found in Supplementary Table S9).

**Table 1.** KEGG pathways enriched among genes that have differential umCpG_rr_ and 5hmC signals and among genes predicted to be targeted by general enhancers with differential hydroxymethylation.

| Differential signal | Genomic annotation | Enriched pathways (FDR) |
| --- | --- | --- |
| 5hmC | gene bodies | Metabolic pathways ($1.3 \times 10^{-11}$) |
| 5hmC | enhancers | Metabolic pathways ($2.3 \times 10^{-4}$) |
| umCpG<sub>rr</sub> | gene bodies | Pentose and glucuronate interconversions (0.0018)<br>Ascorbate and aldarate metabolism (0.071)<br>Metabolic pathways (0.071) |

## DISCUSSION

Optical genome mapping is a powerful tool that complements DNA sequencing. While NGS identifies short InDels and SNPs, OGM unravels complex genomic structures including SVs and CNVs of medium to large sizes. The long reads in OGM enable *de novo* construction of a sample’s genomic structure, which can be particularly complex in cancer. Here we present a complete structural comparison between tumors and matched samples in different stages of disease. By identifying changes that are specific to each stage or sub-diagnosis, the method can be used to track disease progression. Our analysis revealed some well-established as well as some novel somatic aberrations of ccRCC (5, 48–51).

Beyond genetic structure, optical *epigenome* mapping can be obtained simultaneously with OGM at (almost) no extra cost, and adds significant information to it. With our epigenetic profiling addition to commercial OGM, we were able to detect epigenetic changes at multiple levels: global (bulk), locus-specific (population average), and single-molecule (single-cell-like). On the bulk level, we observed a significant reduction of umCpG_rr_ and 5hmC signals in the tumor sample compared with the matched adjacent tissue. This reduction suggests that the tumor sample is both hyper-methylated and hypo-hydroxymethylated, in line with previous studies (8, 15). At the locus-specific level, we found a correlation between gene expression and the umCpG_rr_ signal around transcription start sites and 5hmC signals in gene bodies. Such correlation may imply that gene expression information may be deduced from the same experiment, reducing the need for dedicated RNA sequencing, and adding additional Omics aspects to the same data. Optical epigenome mapping effectively provides single-cell-level information of the locus covered by the method’s long reads. Previously, we showed that the simultaneous recording of the methylation status of gene promoters and their distant enhancers enables generation of epigenetic signatures, and used these signatures to de-convolve cell mixtures (25). In the current study, the methylation and hydroxymethylation state in all DNA molecules were recorded, providing information on the cell-to-cell epigenetic variability. This single-cell-like comparison between tumor and matched tissues identified significantly distinct regions. Some of these regions fall within annotated elements known to be associated with ccRCC (e.g., *PRCC, BAP1, VHL*,*PBRM1*; (3, 37)), further validating the results, and some within coding or non-coding regions not previously linked to ccRCC. Among annotated differentially modified regions, as genes and enhancers, we found significant enrichment of metabolic pathways, and other pathways involving metabolites. Metabolism plays an important role in renal health and disease, and especially in renal cancers (61). As seen here, epigenetic dysregulation of ccRCC-related genes as well as of other genomic regions is highly distinct and may inspire new studies on the mechanism of the disease.

To conclude, this study, using whole-exome sequencing, OGM and optical epigenome mapping, provides a uniquely comprehensive analysis of the spectrum of somatic alteration affecting ccRCC, and demonstrates the potential of combining these methods to understand carcinogenesis. The methodology can be applied to other questions in comparative genomics. Nevertheless, clinical insights will require larger cohort studies.

## Data Availability

The raw data underlying this article are available upon request.

## Supplementary Data

Supplementary Data are provided.

