## Supplementary Data for "Optical Genome and Epigenome Mapping of Clear Cell Renal Cell Carcinoma"

1 Department of Chemistry, Raymond and Beverly Sackler Faculty of Exact Sciences, Tel Aviv University, 6997801 Tel Aviv, Israel; 2 Institute of Experimental Botany of the Czech Academy of Sciences, Olomouc, Czech Republic; 3 Institute of Organic Chemistry, RWTH Aachen University, D-52056 Aachen, Germany; 4 Pediatric Stem Cell Research Institute, Edmond and Lily Safra Children's Hospital, Sheba Medical Center, 52621 Ramat Gan, Israel; 5 Pediatric Nephrology Unit, The Edmond and Lily Safra Children's Hospital, Sheba Medical Center, 52621 Ramat Gan, Israel; 6 Sackler Faculty of Medicine, Tel-Aviv University, 6997801 Tel Aviv, Israel; 7 Susanne Levy Gertner Oncogenetics Unit, The Danek Gertner Institute of Human Genetics, Sheba Medical Center, 52621 Ramat Gan, Israel; 8 Department of Biomedical Engineering, Tel Aviv University, 6997801 Tel Aviv, Israel.

### Supplementary Data - methods and results

#### 1. Whole-exome sequencing

##### a. Library preparation

500 ng DNA were used for library construction. Sequencing libraries were generated using Agilent SureSelect Human All Exon kit (Agilent Technologies) following manufacturer's recommendations and index codes were added to attribute sequences to each sample. DNA samples were sonicated by hydrodynamic shearing system (Covaris) to generate 180-280 bp fragments. Remaining DNA overhangs were converted into blunt ends *via* exonuclease/polymerase activities, and the enzymes were then removed. After adenylation of 3' ends of DNA fragments, adapter oligonucleotides were ligated. DNA fragments with ligated adapter molecules on both ends were selectively enriched using PCR. After PCR, the library was hybridized in the liquid phase with biotin-labeled probes, after which streptavidin-coated magnetic beads were used to capture the exons of genes. Captured libraries were enriched by PCR to add index tags to prepare for hybridization. Resulting products were then purified using the AMPure XP system (Beckman Coulter) and quantified using the Agilent high sensitivity DNA assay on the Agilent Bioanalyzer 2100 system. The qualified libraries were sequenced using Illumina NovaSeq6000 after pooling according to its effective concentration and expected data volume.

##### b. Analysis

Filtering raw sequencing reads was performed using Trim Galore software (v0.6.7; 10.5281/zenodo.5127898), parameters: -q 25 –phred33 –length 36 -e 0.1 –stringency 3 –fastqqc.

Alignment to hg38 was performed with Bowtie2 software (v2.2.5; (1)), parameters: default.

For variant calling, a pipeline by Genome Analysis Toolkit (GATK, v4.2.2.0; (2)) was followed, using Picard (v2.25.4; https://github.com/broadinstitute/picard/) and Samtools (v0.1.19; (3)). Steps:

gatk SortSam; gatk MarkDuplicates; gatk FixMateInformation; gatk BaseRecalibrator --known-sites dbsnp_146.hg38.vcf --known-sites Mills_and_1000G_gold_standard.indels.hg38.vcf; gatk

ApplyBQSR; gatk ValidateSamFile; gatk HaplotypeCaller -ERC GVCF; gatk CombineGVCFs; gatk GenotypeGVCFs; gatk SelectVariants -select-type <SNP | INDEL>; gatk VariantFiltration --filter-expression "QD < 2.0 || MQ < 40.0 || FS > 60.0 || SOR > 3.0 || MQRankSum < -12.5 || ReadPosRankSum < -8.0"; VCFtools (v0.1.16; (4)), parameters: --recode –bed exons.bed.
SnpSift (v4.3t; (5)) program was used to add NCBI dbSNP (v146; (6)) information, and SnpEff (v4.3t; (5)) program was used to annotate the variants and determine the effect of each variant.

#### c. Results – SNPs and short InDels

In the early stage sample pair, the whole-exome sequencing analysis revealed 84,656 variants in the non-tumorous adjacent tissue, and 75,682 in the tumor tissue; 76,177 and 68,169 of them were SNPs, and the rest were short InDels. In the advanced stage samples pair, this analysis revealed 80,543 variants in the non-tumorous adjacent tissue, and 84,220 in the tumor tissue; 72,311 and 75,395 of them were SNPs, and the rest were short InDels (Figure S1.A.). The variants were annotated using SnpEff (v4.3t) and their effect and impact were determined. 1,181 of the normal adjacent sample's variants and 1,411 of the tumor's variants were classified as 'high impact variants' in the early stage pair, determining a severe consequence, and 1,239 and 1,253, respectively, in the advanced samples pair. In the early samples pair, 15,476 protein-coding genes harbored variants in the normal adjacent tissue, and 15,006 in the tumor tissue. Out of them, 14,380 are common for both samples. In the advanced samples pair, 15,012 protein-coding genes harbored variants in the normal adjacent tissue, and 15,235 in the tumor tissue. Out of them, 14,331 are common for both samples (Figure S1.B.). Some of the variants discovered were in genes that are known to be associated with ccRCC (7–9). For example, the *VHL* gene exhibited 1bp InDels classified as high impact in the tumor samples of both sample pairs. Additional SNPs (low depth) were also observed in this gene in both the adjacent tissue and the tumor of the advanced stage pair. In the normal adjacent sample of the early stage pair, the *VHL* gene is wild type. The gene *PBRM1* exhibited several variants in all four samples, including one high impact variant in the tumor sample of the early stage. The genes *SETD2* and *PRCC* displayed several variants in both the advanced stage samples. A list of all genes harboring high impact variants in these samples can be found in supplementary table S1.

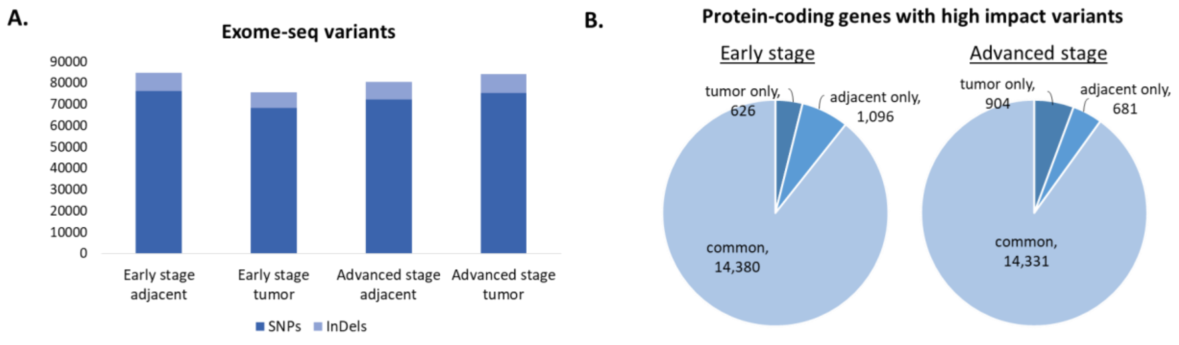


**Figure S1.** Variants discovered by whole-exome sequencing. **A)** Distribution of SNPs and InDels in the tumor and adjacent tissues of the early and advanced stage ccRCC patients. **B)** Protein-coding genes with a high impact variant, signifying severe consequence, in the two sample pairs.

### 2. Synthesis of AdoYnATTO643

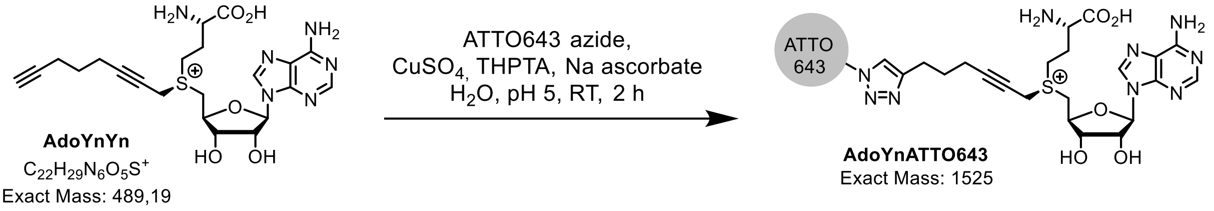


An aqueous solution (400 µL) containing 1.04 µmol AdoYnYn (10), 0.87 µmol ATTO643 azide (ATTO-TEC, $M^+$ = 1036), 25 µmol sodium ascorbate, 0.44 µmol $CuSO_4$ (1 e) and 0.44 µmol THPTA was prepared and the pH was adjusted to 5 with diluted sulfuric acid. The reaction mixture was protected from light and stirred at room temperature. The progress of the reaction was monitored by analytical reversed phase HPLC ($t_R$ = 26.1 min, Prontosil C-18 AQ column, 250 x 4.6 mm, 5 µm, 120 Å, equipped with a C-18 precolumn, 8.0 mm x 4.0 mm, 5 µm, 120 Å) using an acetonitrile gradient (11.9–16.1% in 10 min and 16.1–70% in 20 min) in aqueous TFA solution (0.01 % TFA) with a flow of 1 mL/min and detection at 260 nm and 640 nm. The reaction was terminated by freezing the solution after 2 h and the product purified by preparative reversed phase HPLC ($t_R$ = 25.0 min, Prontosil C-18 AQ column, 250 x 20 mm, 5 µm, 120 Å, equipped with a C-18 precolumn, 30 x 20 mm, 5 µm, 120 Å) using an acetonitrile gradient (11.9–16.1% in 10 min and 16.1-70% in 20 min) in aqueous TFA solution (0.01 % TFA) with a flow of 10 mL/min and detection at 260 nm and 640 nm. The product was collected and the volume was reduced by lyophilization. The final concentration of the fluorescent AdoMet analogue AdoYnATTO643 was obtained using UV-Vis spectroscopy ($\varepsilon_{643}$ = 150,000 L $mol^{-1}$ $cm^{-1}$). The obtained yield was 55.9%. Mass spectrum was acquired with a Thermo Fisher Scientific LTQ Orbitrap XL electrospray ionization mass spectrometer (ESI-MS) in positive ion mode (70 eV; Supplementary Figure S2). m/z (%) = 762.84 $[M]^{2+}$ (100).

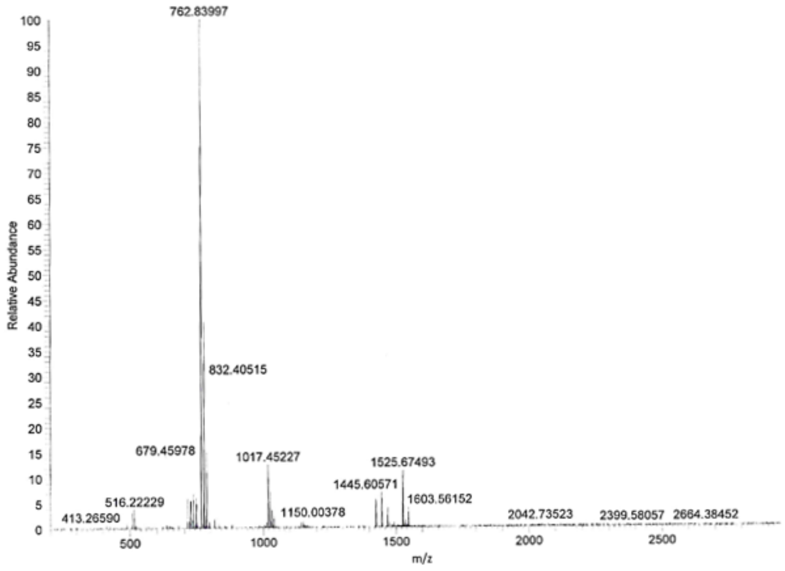


**Figure S2.** ESI mass spectrum in positive ion mode of AdoYnATTO643. The product (100%) has m/z of 762.84 $[M]^{2+}$.

### 3. Synthesis of DBCO-ATTO643

ATTO-643 modified with carboxylic acid (ATTO-643-COOH, 5 mg, 5.3 µmol, ATTO-Tec) was dissolved in 1 mL dry DMF under argon. DIPEA (3 µL, 17.2 µmol) and HATU (2.5 mg, 6.6 µmol) were added to the solution and stirred at 0°C for 2 hours. Dibenzocyclooctyne-amine (1.5 mg, 5.4 µmol, Sigma Aldrich) was added, and the solution was stirred at 0°C for 2 hours and monitored using analytical reverse-phase HPLC (Prontosil C-18 AQ column, 250 x 4.6 mm, 5 µm, 120 Å; mobile phase: acetonitrile in $H_2O$ containing 0.1% TFA (v/v); gradient from 10% to 100%; flow rate: 1 mL/min). Upon completion, purification of the product was carried out by preparative reverse-phase HPLC (Prontosil C-18 AQ column, 250 x 20 mm, 5 µm, 120 Å; mobile phase: acetonitrile in $H_2O$ containing 0.1% TFA (v/v); gradient from 10% to 100%; flow rate: 5 mL/min) and yielded DBCO-ATTO643 (3.9 mg, 61%) as a blue powder.

### 4. Genome-wide structural variations in the early and advanced stage sample pairs

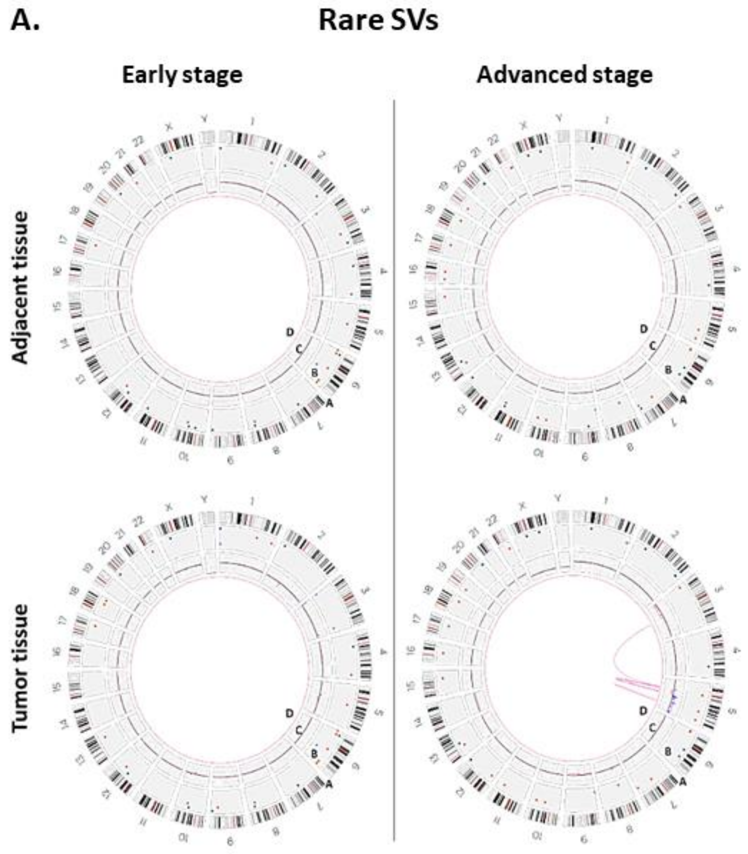


**Figure S3.** Circos plot view of Variant annotation pipeline results for early and advanced stage samples. **A)** Rare SVs, not present in Bionano Genomics' control database. The circos plots display cytobands ("A") including centromeres (bands highlighted in red), all structural variations as colored dots – insertions (dark green, 1st level in "B"), deletions (orange, 2nd level in "B"), inversions (light blue , 3rd level in "B"), duplications (light purple, 4th level in "B") and translocations ("D"), as well as copy number variations ("C") with a baseline set to two copies, where DNA gains (blue peaks) and DNA losses (red peaks) are visualized. The outermost numerical track corresponds to chromosome number.

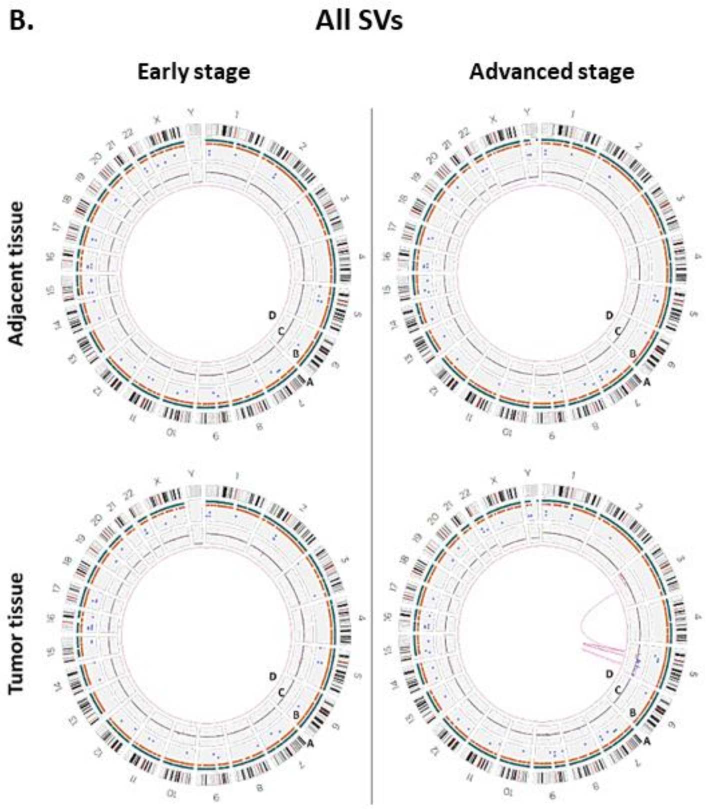


**Figure S3. B)** All detected SVs in the two sample pairs.

### 5. SVs overlapping ccRCC-relevant genes

The gene *PBX1* (Pre-B-cell leukemia homeobox transcription factor 1) overlaps a deletion unique to the early stage tumor sample. This gene is localized on the long arm of chromosome 1 and was shown to have an oncogenic role in the progression of ccRCC (11, 12). Wei and co-authors concluded that upregulation of *PBX1* efficiently promoted ccRCC cell proliferation, and that *PBX1* knockdown inhibits ccRCC cell viability, proliferation and cell cycle progression *in vivo* and *in vitro* (12). The *EPHB1* gene, overlapping a deletion unique to the normal adjacent sample of this pair, is localized on the long arm of chromosome 3 and encoding Eph receptor tyrosine kinase. Protein tyrosine kinase genes are the largest family of oncogenes (13). However, Zhou *et al.* revealed decreased expression of *EphB1* in all RCC carcinomas including ccRCC, suggesting that it may play a tumor-suppressive role in cancer (14). In the advanced stage tumor, genes encoding for microRNAs, which are known to dysregulate expression in cancer states (15), uniquely overlap with the detected SVs. Additionally, a 2.2 Mbp deletion in chromosome 5, detected in this sample, overlaps several genes including *ESM1* (endothelial cell specific molecule-1), which was previously found to be markedly overexpressed in ccRCC (16). However, in the case of a deletion covering the gene, overexpression is not likely. Other genes overlapping SVs (see full list in Supplementary Table S4) were not associated with ccRCC before.

### 6. Copy number variation profiles of early and advanced stage sample pairs

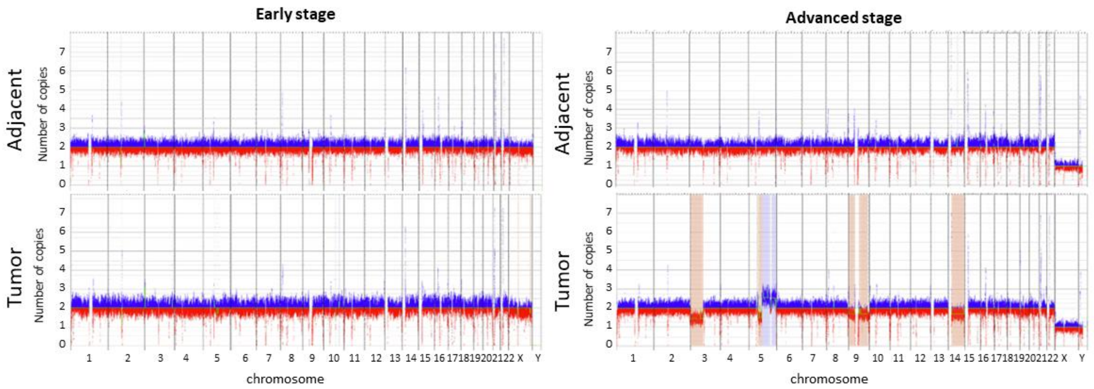


**Figure S4.** Copy number variation profiles of early and advanced stage sample pairs. The Copy number analysis is based on molecule-to-reference alignment and coverage normalization. Y axes display number of copies with a baseline set to two copies. DNA gains are colored in blue and DNA losses are colored in red. Aneuploidies are highlighted with vertical boxes in matching colors.

### 7. Gene fusion events

Fusion genes are chimeric genes formed from concatenation of two or more independent genes as a byproduct of genomic instability in cancer. Hence, gene fusions can contribute to disruption of gene expression stability caused by expression of genes from novel promoters or by fusion-initiated change of functional products dosage (17, 18). A putative gene fusion event detected in the early stage ccRCC tumor involves the genes *SLC35E2B* and *CDK11A.* It was identified in the 1p36.33 cytoband localized on the distal part of the short arm of chromosome 1. *CDK11A* codes for cyclin dependent kinase 11A, which is known to be essential for eukaryotic cell cycle regulation or a candidates for tumor suppressors that modulate regulatory pathways critical in development and disease (19, 20). Another putative gene fusion, detected in both early stage samples, connects the genes *LINC01266* and *CNTN6* on the 3p26.3 cytoband localized on the distal part of the short arm of chromosome 3. *LINC01266* encodes for long intergenic non-protein coding RNA (lncRNA) 1266. lncRNAs play an important role in renal cancer (21, 22). The 3p26.3 cytoband contains three consecutive genes encoding closely related neuronal immunoglobulin cell adhesion molecules, *Contactin-6* (*CNTN6*) is one of them (23). This cytoband is also associated with deletions that can cause 3p-deletion syndrome (Del3p), typically characterized by growth retardation, developmental delay or renal and gastrointestinal abnormalities. Another putative fusion event recognized in the early stage pair involves the genes *SLC35E2B* and *CDK11A*. This fusion, as well as the advanced stage fusion of *PCDHA* cluster and *AC011346*, affect genes that are known to be essential for eukaryotic cell cycle regulation or tumor suppressor candidates that modulate regulatory pathways critical in development and disease (19, 20). A putative gene fusion of *SPIDR* (scaffold protein involved in DNA repair) and *PRKDC* (protein kinase, DNA-activated, catalytic subunit) in both advanced stage adjacent tissue and tumor, is localized on the short arm of chromosome 8 in 8q11.21 cytoband. These two genes are separated by the gene *CEBPD* (CCAAT enhancer binding protein delta). The fusion is a result of a duplication of a region involving the last nine exons of *SPIDR*, *CEBPD* and the last 16 exons of *PRKDC*. This fusion can lead to incorrect *PRKDC* gene product. PRKDC (also known as DNA-PKcs) is a member of the protein kinases family (PIKK) that plays an essential role in response to DNA double-strand damage (24) and in cancer metastasis (25), as overexpression of *PRKDC* leads to a maintenance of tumor balance and helps the tumor cells resist treatment with DNA damage, and as the absence of DNA-PKcs results in a secretion of anti-metastatic factors, thus inhibiting cell migration and invasion (25).

### 8. Genome-wide profiles of 5hmC and unmodified CpGs

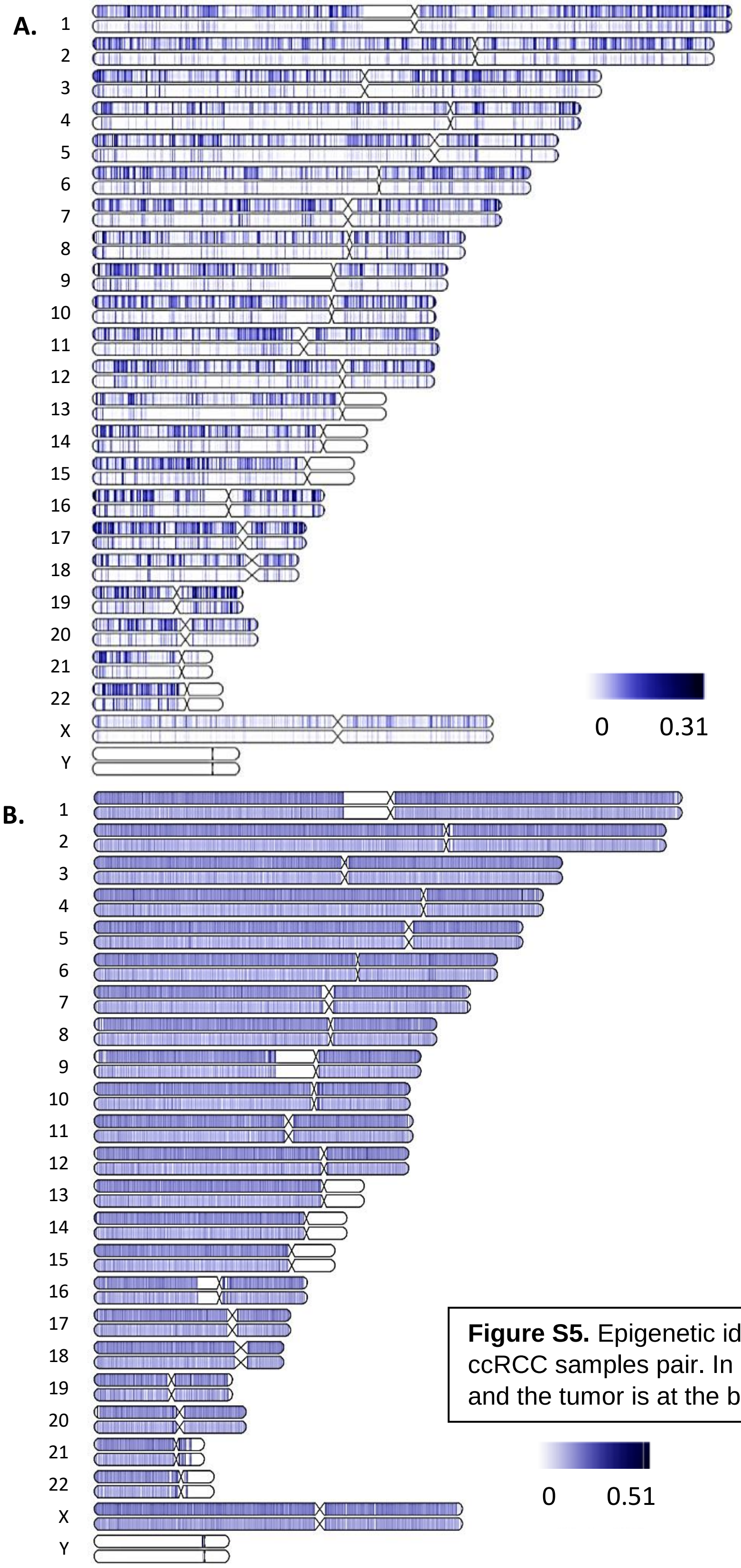


**Figure S5.** Epigenetic ideograms of all chromosomes in the early stage ccRCC samples pair. In both ideograms, the adjacent sample is on top and the tumor is at the bottom. **A)** 5hmC profile. **B)** umCpG$_{rr}$ profile.

### 9. Fluorescence microscopy image of single molecules with umCpG$_{rr}$ repeats

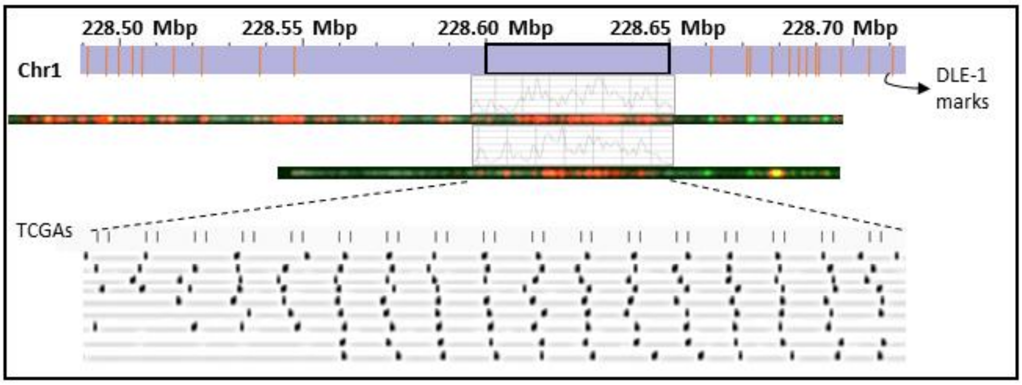


**Figure S6.** Fluorescence microscopy images of two molecules containing a repeat array aligned to chromosome 1, displayed in Figure 3.C. Red dots are umCpG$_{rr}$ signals, depicted in the digitized scheme as black dots. The green dots are genetic barcode labels, corresponding to DLE-1 marks. Above the fluorescent repeat array there is a corresponding intensity plot of the red channel.

### 10. umCpG$_{rr}$ scores correlate with gene expression and with WGBS scores in the GM12878 cell line

Single-molecule umCpG$_{rr}$ maps of three replicates of the B-lymphocyte cell line GM12878 were adapted from Sharim et al. (26) and merged. Publicly available Whole-genome bisulfite sequencing (WGBS) data of the cell line was obtained from GEO accession GSM2308633. In order to match the two signals, the umCpG score was calculates from the WGBS data (1-mCpG score), and the average of the two replicates was calculated. Gene expression data from RNA sequencing for GM12878 was obtained from GEO accession GSE88583, and the average TPM score of two replicates was calculated. Protein coding genes were divided into four groups based on their average TPM score. Unexpressed genes were defined as genes with TPM value <= 0.01 (~4200 genes). The other expression groups are three equal quantiles of the expressed protein-coding genes (~4700 genes in each group). Mean umCpG($_{/rr}$) signals along genes were calculated using DeepTools *computeMatrix* (v3.4.1, (27)) in scale-regions mode, where each gene was scaled to 15 kbp and divided into 300 bp bins. Optical epigenome mapping signals and WGBS signals were normalized between 0 and 1 for comparison. Figure S7 shows a reduction in umCpG($_{/rr}$) singal in gene bodies, as gene expression increases (S7.B.), as well as an increase in umCpG($_{/rr}$) singal around the gene's TSS as gene expression increases (S7.C.).

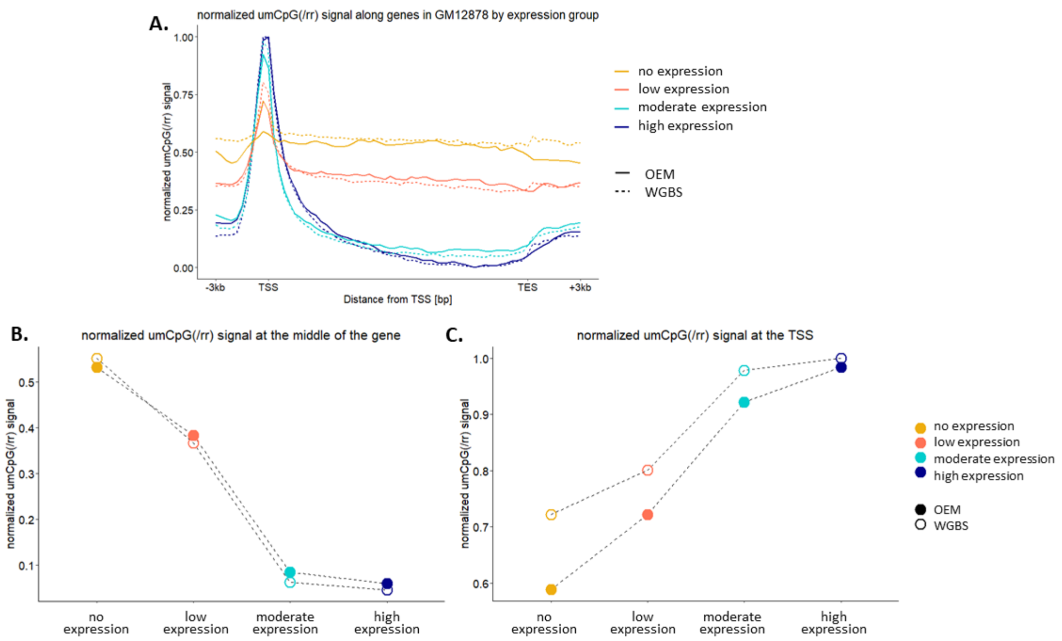


**Figure S7. A)** Mean normalized umCpG($_{/rr}$) score in optical epigenome mapping (OEM) and whole genome bisulfite sequencing (WGBS) in the human cell line GM12878, in groups of genes divided according to their expression. **B)** Mean normalized umCpG($_{/rr}$) score in gene midpoints. **C)** Mean normalized umCpG($_{/rr}$) score at the TSS.

### 11. umCpG$_{rr}$ and 5hmC levels around ccRCC-related enhancers in the early stage samples

Both umCpG$_{rr}$ and 5hmC signals peak at enhancer midpoints. The signal of both modifications is reduced in the tumor sample compared with the normal adjacent tissue.

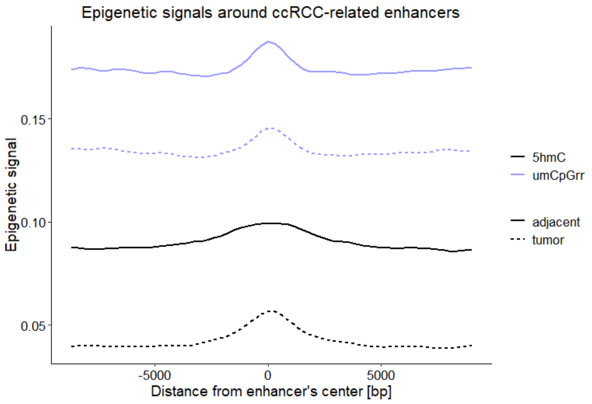


**Figure S8.** umCpG$_{rr}$ (violet) and 5hmC (black) signals of the early stage adjacent (solid line) and tumor (dashed line) samples around ccRCC-related enhancers.

### 12. Differentially modified genomic windows

1,114,617 non-overlapping 1 kbp genomic windows that contain at least one umCpG$_{rr}$ recognition site were covered in both samples by at least 20 molecules (16 of these windows were removed due to 0 in the t-test denominator). For 5hmC, 2,744,450 non-overlapping 1 kbp genomic windows that contain at least one CpG site were covered in both samples by at least 20 molecules (242,338 of these windows were removed due to 0 in the t-test denominator). In the following figure, the fold change between the adjacent and tumor samples (early stage) is plotted as a function of the FDR-adjusted p-value (q-value) of these windows. The dynamic range of $umCpG_{rr}$ scores is higher than that of the 5hmC score, as 5hmC is a scarce modification (28), resulting in higher relative variability in 5hmC. Accordingly, the q-value scores are less significant in the 5hmC windows, even though the fold change values are more dramatic.

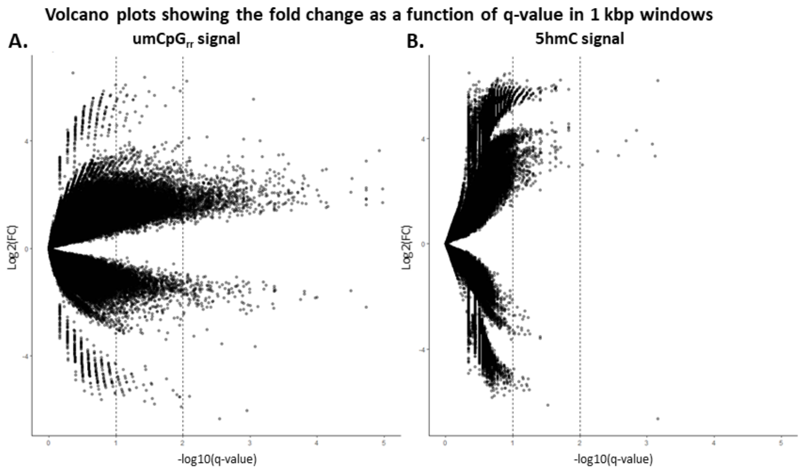


**Figure S9.** 1 kbp genomic windows, spread according to their adjusted p-value (q-value) in the differential modification test, and their fold change (adjacent/tumor), in the early stage samples pair. **A)** $umCpG_{rr}$ score. **B)** 5hmC score.

### 13. Legend for Supplementary tables S1-S9 (in the attached .xlsx file)

**S1.** Genes with high impact variants (SNPs/InDels) in the early and advanced stage ccRCC tumors and adjacent tissues.

**S2.** Statistics of *de novo* assemblies for early and advanced stage sample pairs.

**S3.** Bionano Genomics' Variant Annotation Pipeline (VAP) results, for early and advanced stage sample pairs.

**S4.** List of genes overlapping with SVs identified by VAP for early and advanced stage sample pairs.

**S5.** List of genomic regions with copy number variations (CNVs), detected by Bionano Genomics' CNV analysis, in the early and advanced stage sample pairs.

**S6.** Optical epigenome mapping statistics.

**S7.** List of differentially modified ($umCpG_{rr}$ and 5hmC) genes and promoters.

**S8.** List of differentially modified ($umCpG_{rr}$ and 5hmC) enhancers.

**S9.** List of enriched biological terms among differentially modified ($umCpG_{rr}$ and 5hmC) elements.
